# Evolutionary states and trajectories characterized by distinct pathways stratify ovarian high-grade serous carcinoma patients

**DOI:** 10.1101/2022.08.30.505808

**Authors:** Alexandra Lahtinen, Kari Lavikka, Yilin Li, Anni Virtanen, Sanaz Jamalzadeh, Kaisa Huhtinen, Olli Carpén, Sakari Hietanen, Antti Häkkinen, Johanna Hynninen, Jaana Oikkonen, Sampsa Hautaniemi

**Affiliations:** Research Program in Systems Oncology, Research Programs Unit, Faculty of Medicine, University of Helsinki, Helsinki, Finland; Department of Pathology, University of Helsinki and HUS Diagnostic Center, Helsinki University Hospital, Helsinki, Finland; Cancer Research Unit, Institute of Biomedicine and FICAN West Cancer Centre, University of Turku, Turku, Finland; Department of Pathology, University of Helsinki and HUSLAB, Helsinki University Hospital, Helsinki, Finland; Department of Obstetrics and Gynaecology, University of Turku and Turku University Hospital, Turku, Finland

**Author notes:** Corresponding authors’ (J.O.); (S.Ha.).

## Abstract

Ovarian high-grade serous carcinoma (HGSC) is typically diagnosed at an advanced stage, with multiple genetically heterogeneous clones existing in the tumors long before therapeutic intervention. Herein we characterized HGSC evolutionary states using whole-genome sequencing data from 214 samples of 55 HGSC patients in the prospective, longitudinal, multiregion DECIDER study. Comparison of the tissues revealed that site-of-origin samples have 70% more unique clones than the metastatic tumors or ascites. By integrating clonal composition and topology of HGSC tumors we discovered three evolutionary states that represent a continuum from genomically highly variable to stable tumors with significant association to treatment response. The states and their evolutionary trajectories were characterized by unique, targetable pathways, which were validated with RNA-seq data. Our study reveals that genomic heterogeneity is unaffected by the current standard-of-care and pinpoints effective treatment targets for each group. All genomics data are available via an interactive visualization platform for rapid exploration.

## INTRODUCTION

High-grade serous carcinoma (HGSC) is the most common epithelial ovarian cancer subtype. HGSC has median latency of more than 10 years (Gerstung et al., 2020) and is typically diagnosed at the metastasized state. Accordingly, treatment-naïve HGSC tumors have undergone expansions of genomically distinct clones leading to a highly heterogeneous disease (Bashashati et al., 2013; Geistlinger et al., 2020; Labidi-Galy et al., 2017; Masoodi et al., 2020), and already contain drivers for tumor progression and therapy resistance (Castellarin et al., 2013; Kozłowska et al., 2018; McPherson et al., 2016; Nath et al., 2021; Schwarz et al., 2015). Thus, while 80% of the HGSC patients respond well to the current standard-of-care, almost all patients experience relapse within a few years after the diagnosis, leading to a five-year survival of only 40% (Torre et al., 2018). We hypothesized that integrating genomic heterogeneity to clonal compositions from phylogenetic analysis enable stratification of the HGSC patients into tumor evolutionally defined groups characterized by unique signaling cascades.

“Multi-layer Data to Improve Diagnosis, Predict Therapy Resistance and Suggest Targeted Therapies in HGSOC” (DECIDER; ClinicalTrials.gov identifier: NCT04846933) is a prospective, longitudinal, multiregion observational study that began recruitment in 2012. In the DECIDER trial multiregion samples are collected from the consented stage III-IV HGSC patients and subjected to whole-genome sequencing (WGS) and RNA-sequencing. A goal of DECIDER is to characterize intra- and inter-tumor heterogeneity in HGSC patients and test whether heterogeneity is associated with treatment response. Here, we integrated clonal compositions and phylogenetic topologies using whole-genome sequencing data of 214 diagnostic and post-treatment samples from 55 HGSC patients, to investigate evolutionary states, trajectories and pathways characterizing them. Transcriptomics data from DNA-matched samples were used to further analyze the changes in the pathway activities at the course of treatment. As the detected oncogenic pathways include several drug targets, our efforts facilitate identifying robust patient groups, as well as personalized treatment modalities.

## RESULTS

### Genomics profiling of tumor biopsies in the DECIDER cohort

We prospectively enrolled 55 HGSC patients and collected 214 samples from tubo-ovarian (ovaries and fallopian tubes) tumors, intra-abdominal metastases (omentum, peritoneum, and bowel mesenterium), other tissues (lymph nodes, liver), and ascites before and after treatments (Figure 1A, Table 1 and Table S1). The patients were treated according to the HGSC standard-of-care guidelines (Colombo et al., 2019), *i.e*., surgery and platinum and taxane combination chemotherapy with angiogenesis and/or poly (ADP-ribose) polymerase (PARP) inhibitors as maintenance therapy (Figure 1B). The clinical information including the treatments for the patients in the cohort is available in Table S2. Archived hematoxylin and eosin (H&E) stained and immunohistochemical slides were re-evaluated by a pathologist to verify histologic diagnosis of HGSC for each patient according to the current diagnostic criteria (Editorial Board, 2020).

**Figure 1.**
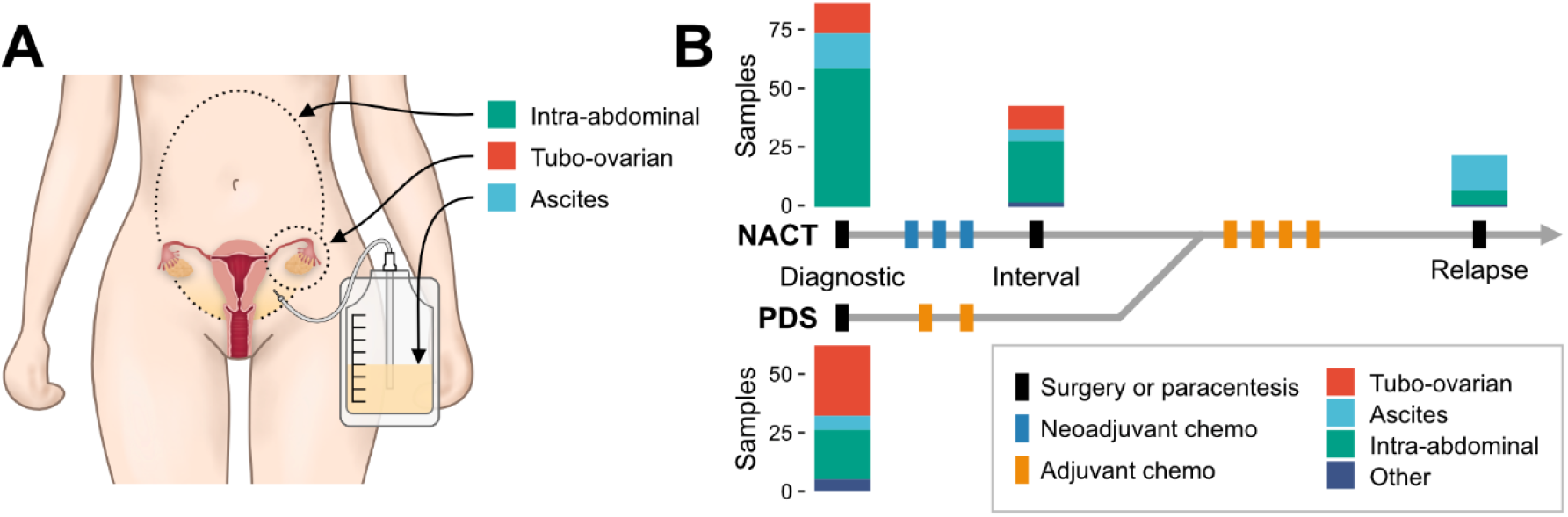
Longitudinal multi-tissue HGSC cohort enables construction of phylogenetic trees and assessment of clonal compositions at subclonal resolution. **A.** Anatomical sites for samples. Intra-abdominal sites include omentum, peritoneum, and bowel mesenterium. Tubo-ovarian sites include fallopian tubes and ovaries. Other metastatic tissue sites include lymph nodes and liver. **B.** Sample collection at the time diagnosis, at interval surgery, and at the time of disease progression. PDS, primary debulking surgery; NACT, neo-adjuvant chemotherapy.

**Table 1.** Characteristics of 55 patients included in the study

| Characteristic | Group | Total |
| --- | --- | --- |
| Age group |  |  |
|  | < 70 | 29 (52.7%) |
|  | ≥ 70 | 26 (47.3%) |
| Treatment strategy |  |  |
|  | PDS | 19 (34.5%) |
|  | NACT | 36 (65.5%) |
| Stage |  |  |
|  | IIIB | 1 (1.8%) |
|  | IIIC | 34 (61.8%) |
|  | IVA | 11 (20.0%) |
|  | IVB | 9 (16.4%) |
| Primary therapy outcome |  |  |
|  | Complete | 25 (45.5%) |
|  | Partial | 22 (40.0%) |
|  | Progressive | 6 (10.9%) |
|  | Other | 2 (3.6%) |
| PFI, months |  |  |
|  | <6 | 22 (40.0%) |
|  | 6-12 | 11 (20.0%) |
|  | 12-24 | 14 (25.5%) |
|  | >24 | 5 (9.0%) |
|  | NE | 3 (5.5%) |
| OS, months |  |  |
|  | <24 | 17(31.0%) |
|  | 24-36 | 19 (34.5%) |
|  | >36 | 19 (34.5%) |
| Progression |  |  |
|  | yes | 52 (94.5%) |
|  | no | 3 (5.5%) |

PDS, primary debulking surgery; NACT, neoadjuvant chemotherapy; PFI, platinum free interval; NE, not evaluable follow-up assessments; OS, overall survival.

Samples were whole-genome sequenced (36x median coverage, range 24-96x) and processed in the Anduril bioinformatics workflow platform (Cervera et al., 2019), as described in Methods. On average, we detected 15,102 mutations per patient (range 5,451-53,458). Mutational patterns varied in the patients, except for *TP53*, which was mutated in all patients (Table S3). The most significantly mutated genes (*p* < 0.001) after *TP53* were *BRCA2* (11%), *AGAP6* (7%), *MGA* (9%), and *RB1* (5%). The most frequently amplified copy-number segments at 3q26.2, 3q29, 8q24.21, and 8q24.3 loci were detected in more than half of the patients (Table S4). The top five frequently amplified genes within the COSMIC Cancer Census genes (Sondka et al., 2018) were *MYC* (8q24.21, 36%), *RECQL4* (8q24.3, 31%), *MECOM* (3q26.2, 22%), *CCNE1* (19q12, 22%), and *LYL1* (19p13.13, 18%). The regions deleted in more than half of the patients included 1p36.11, 2q22.1, 3p21.31, 7p22.1, 10q24.32, 11q23.3, 12q24.33, and 14q32.33 (Table S4). The most frequently deleted genes within the COSMIC Cancer Census genes were *CDH1* (15q21.1, 13%), *FANCA* (16q24.3, 11%), and *CDKN2A* (9p21.3, 11%).

To provide interactive exploration of our cohort, all genomics and key clinical data are available via an interactive visualization tool GenomeSpy at https://csbi.ltdk.helsinki.fi/p/lahtinen_et_al_2022/. GenomeSpy enables rapid navigation of genomic data, and allows filtering, sorting, grouping, and aggregating the samples, as well as bookmarking the most interesting analyses. This enables anyone to rapidly see frequencies of mutations and copy-number variations (CNVs) of any gene or genomic region, as well as their association with platinum free interval (PFI), whole-genome duplications and other genomic or clinical features. For example, approximately half of HGSCs tumors exhibit homologous recombination deficiency (HRD), a predictive biomarker for response to platinum and PARP inhibitors (R. E. Miller et al., 2020). We quantified HRD-related mutational signatures based on COSMIC v3.1 single base substitution signatures (Alexandrov et al., 2020) and included the signatures in the visualization. Further, *CCNE1* is amplified in 15–20% of HGSC patients and is predictive of HGSC five-year progression free survival (Stronach et al., 2018). The GenomeSpy visualization shows that high *CCNE1* amplifications are associated with low HRD in the DECIDER cohort, supporting the results in (Takaya et al., 2020).

### Phylogenies reveal highly diverse clonal compositions within and between the HGSC patients

Phylogenetic trees and clonal compositions were constructed for all patients using somatic mutations adjusted for CNVs (Methods). The DECIDER cohort contains samples from multiple anatomic sites in various time-points at the patients’ treatment course, which makes visualizing clonal structures challenging with the traditional fishplots (C. A. Miller et al., 2016). To visualize tumor evolution results from longitudinal and multiregion cohorts, we developed a novel visualization concept called ‘jellyfish’ (Supplementary Jellyfish Plots). Jellyfish captures the phylogeny of the subclones and enables the comparison of clonal compositions of samples that are acquired from multiple anatomical sites at the same or different time points.

An example of a jellyfish visualization of patient EOC742, who had partial response to the first-line treatment and overall survival of 38 months, is shown in Figure 2A. The clonal compositions of the diagnostic samples obtained from ascites and omentum include two shared subclones present in almost equal proportions, as well as unique subclones. The ascites sample obtained at the second progression reveals a drastically different clonal composition and two entirely new subclones characterized by the genes *ERCC4* (major clone) and *MYH9* (minor clone). Based on clonal compositions, the dominant clones in the relapse sample originated from the same ancestral clone as major clones in the two diagnostic samples.

**Figure 2.**
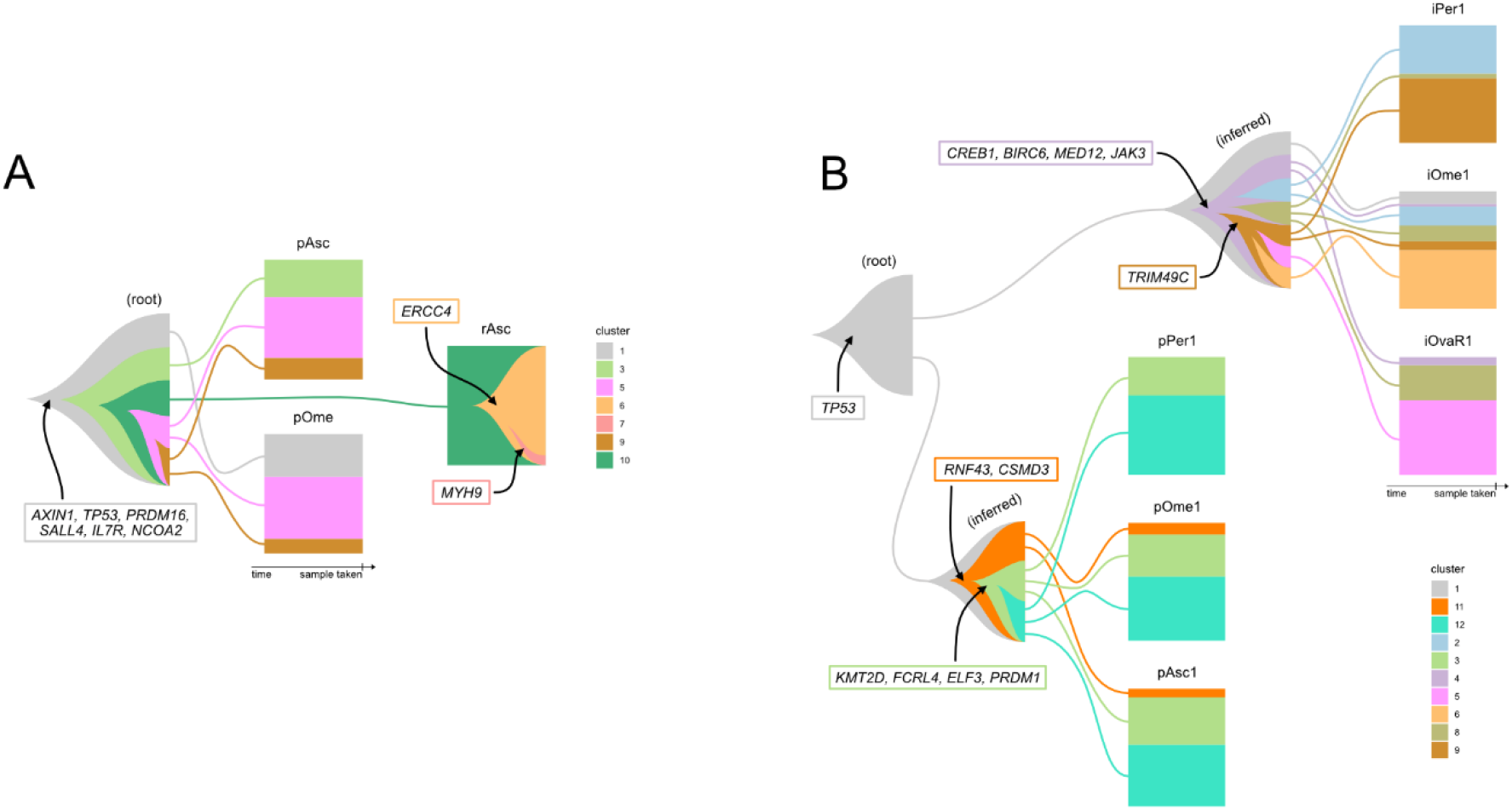
Exemplary jellyfish plots illustrating clonal compositions at various time points. **A.** Jellyfish plot of patient EOC742. The rectangular boxes represent ascites and omental samples collected at the time of diagnosis (pAsc and pOme), and ascites sample at second progression (rAsc). The right edge of each box shows the proportion of subclones in the sample. In the relapse sample, one can see two new subclones observed only at progression (clone 6, clone 7). The inferred sample, root, shown on the left, represents a hypothetical sample with subclones that are inherited by multiple real samples. The lines depict the origins of the clones. **B**. Jellyfish plot of patient EOC153 showing drastic subclonal changes in multiple tissues occurring after NACT treatment.

Overall, clonal compositions revealed high subclonal heterogeneity in the samples prior to the treatments. In line with earlier studies (de Witte et al., 2022; S. Lee et al., 2020; Masoodi et al., 2020; Patch et al., 2015), only a small fraction of patients in the DECIDER cohort showed major changes of the clonal structure or genomics landscape before and after the primary treatment. A rare example of a complete change in clonal composition affecting multiple tissues can be observed in the jellyfish plot of patient EOC153, who had a complete response to the first-line therapy and overall survival 41 months (Figure 2B). The clonal compositions after neoadjuvant chemotherapy (NACT) in all three samples were completely different to the three samples at diagnosis. The clonal compositions of the diagnostic samples were very similar to each other, whereas the NACT treated samples were very dissimilar to each other.

### Heterogeneity is patient and tissue specific

We next quantified heterogeneity at the subclonal level. The two major characteristics of clonal compositions that collectively reflect the heterogeneity measured within and between the samples in a patient are clonal complexity (the number of subclones in the sample) and clonal divergence (the number of unique subclones in the sample). Heterogeneity analyses were carried out in all tissue and ascites samples (*n* = 207).

We first estimated the sample-level clonal complexity, which represents the effective number of clones adjusted for their cellular frequencies (Methods). The sample-level clonal complexity was then averaged to capture the impact of patient, tumor site, and treatment with analysis of variance (ANOVA). Clonal complexity was highly patient specific (*p* = 2.1 · 10^-6^) and could not be explained by the anatomical site (*p* = 1.5 · 10^-3^), as shown in Table S5. Tubo-ovarian samples exhibited significantly more clonal complexity than the other tissue sites (*p* = 0.02) unlike intra-abdominal metastases and ascites (*p* = 0.06 - 0.87). Sample-level clonal complexity did not change significantly before and after the treatments (*p* = 0.72).

To quantify sample-level clonal divergence, we first estimated the number of subclones unique to a sample with respect to the global subclone distribution in all samples of the patient. Clonal divergence was averaged to capture and evaluate the impact of patient, tumor-site, and treatment phase-specificity with ANOVA (Table S5). Like clonal complexity, clonal divergence was highly patient-specific (*p* = 2.2 · 10^-16^). Interestingly, tubo-ovarian tumors featured 70% more unique clones than the other tumor sites (*p* = 5.4 · 10^-8^), whereas ascites samples exhibited 14% less unique clones than the other sites (*p* = 6.6 · 10^-4^). Sample-level clonal divergence did not change significantly before and after the treatments (*p* = 0.39).

### Quantification of tumor heterogeneity enables clustering of HGSC patients into evolutionary states with a distinctive temporal order

As HGSC tumors exhibited both highly patient-specific clonal complexity and divergence, we sought to identify evolutionary states with common clonal structure. Since neither clonal complexity nor divergence alone produced distinct states (results not shown), we clustered the patients along both quantities (Figure 3A). The patients clustered into three distinct states (Figure 3B; Table S6; Methods). Cluster 1 consisted of 20 patients and was characterized by a small number of clones (1.79) and high clonal divergence (1.66). An exemplary jellyfish plot of a Cluster 1 patient is illustrated in Figure 3C. Cluster 2 consisted of 16 patients and was characterized by a high number of clones (3.66) and the lowest clonal divergence (1.23). An exemplary jellyfish plot of a Cluster 2 patient (Figure 3D) demonstrates a stable, almost identical clonal composition across all the tissue sites and ascites. Cluster 3 consisted of 18 patients and was characterized by a high number of clones (2.43) and the highest clonal divergence (2.63). An exemplary jellyfish plot of a Cluster 3 patients features an almost unique clonal composition for every sample (Figure 3E).

**Figure 3.**
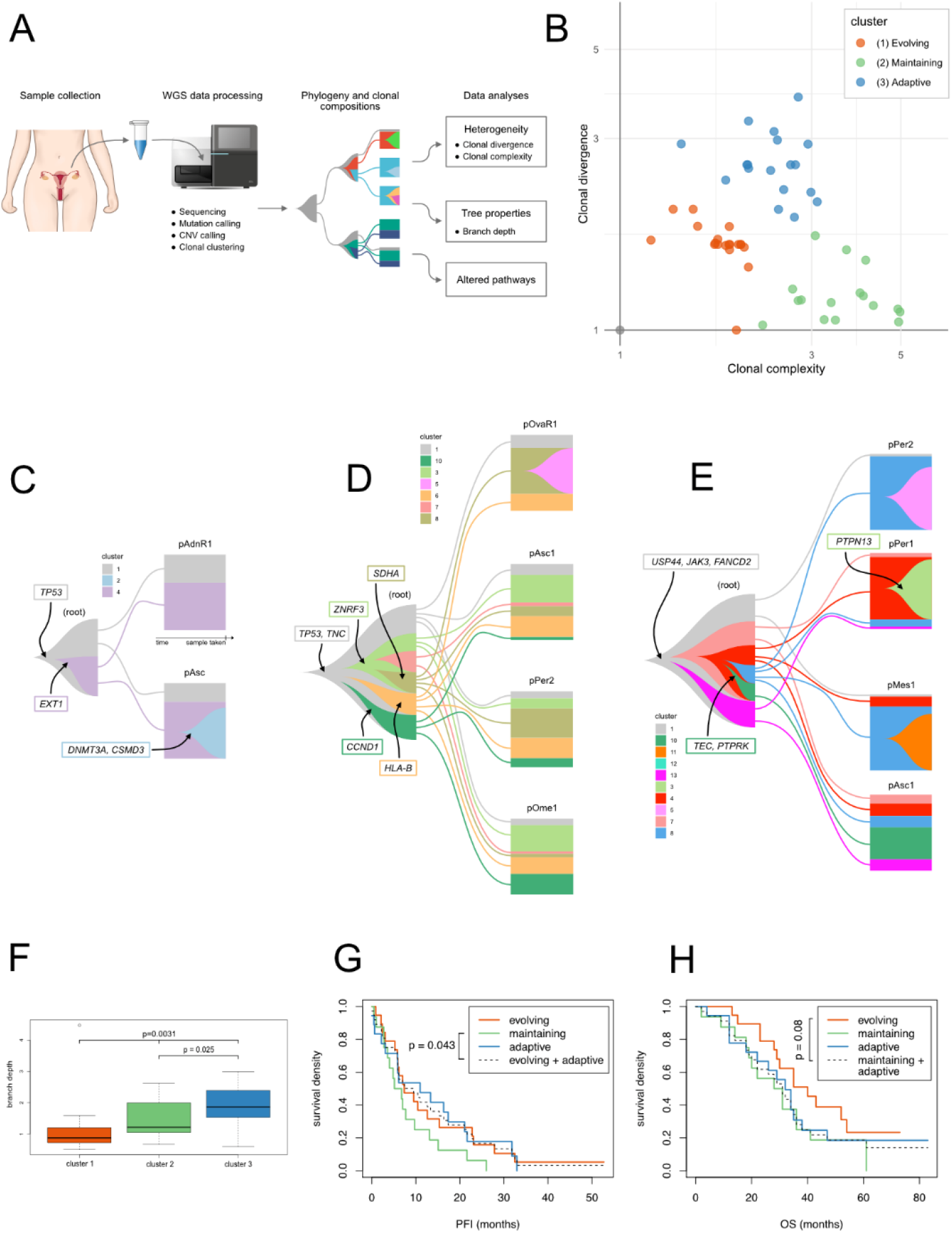
Integration of clonal complexity and divergence enables clustering of HGSC patients into evolutionary states. **A.** The workflow of the study combines three main lines of evolutionary characterization: estimation of evolutionary states based on clonal abundances, temporal order based on the phylogenetic tree topology, and characterization of the underlying biological mechanisms based on pathway aberrations. **B.** The 54 patients (EOC286 was excluded due to the lack of diagnostic samples) clustered into three distinct states along clonal complexity (X-axis) and clonal divergence (Y-axis). The placement of each patient in the clustering is available in the corresponding jellyfish plot (Supplementary Jellyfish Plots). **C.-E**. Jellyfish plots illustrating typical clonal compositions in **C.** Cluster 1 (evolving), patient EOC883, **D**. Cluster 2 (maintaining), patient EOC1129, and **E.** Cluster 3 (adaptive), patient EOC136. These plots show only diagnostic samples and full Jellyfish plots can be seen in Supplementary Jellyfish Plots. **F.** Box-whisker plot of the branch depths of the three evolutionary states. Branch depth corresponds to the clonal pseudo-age. **G-H.** Kaplan-Meier for **G.** platinum-free interval (PFI) and **H.** overall survival (OS) of patients from three evolutionary states.

We tested whether the clonal complexity and divergence were confounded by the number of samples using ANOVA. Clonal complexity and divergence variation explained by clustering were not significantly contributed by the number of samples (complexity: *p* = 0.70; divergence: *p* = 0.46). Furthermore, clustering had a significant explanatory power on clonal complexity (*p* = 1.5 · 10^-12^) and clonal divergence (*p* = 3.6 · 10^-10^) over the number of samples.

To assess the degree of evolvement or clonal pseudo-age of the three states we used the clonal branches along the topology of the phylogenetic trees (Table S6; Methods). Cluster 3 featured the highest clonal pseudo-age, followed by Cluster 2 and Cluster 1 (Figure 3F). The pseudo-ages explained by the clustering were not significantly contributed by the number of samples (*p* = 0.27), and the clustering had a significant explanatory power over the number of samples (*p* = 0.04). Accordingly, as Cluster 1 exhibited the shortest pseudo-age, it is subsequently referred to as ‘evolving state’. Cluster 2 featured a stable clonal composition maintained across all samples and is referred to as ‘maintaining state’. Cluster 3 was distinguished by the unique clonal composition in the samples and is therefore further referred to as ‘adaptive state’.

To investigate whether the evolutionary states were associated with treatment response and survival, we performed survival analysis of patients belonging to the three clusters. The patients from the maintaining state showed shortest times to relapse, determined by platinum-free interval (PFI), and lowest variance, as compared to patients from evolving and adaptive states (*p* = 0.043, Figure 3G). In terms of overall survival (OS), the patients from the evolving state demonstrated the best survival odds (*p* = 0.08, Figure 3H), as compared to patients from maintaining and adaptive states.

### Evolutionary states are characterized by distinct pathways at genomic and transcriptomic levels

As biological mechanisms underlying evolutionary states may open possibilities for targeted therapies, we next proceeded to explore pathways associated with the states. As the three states showed significantly different degrees of clonal pseudo-age, we collected subclonal mutations for each state, annotated them with the genes, and identified enriched biological pathways for each state, as well as pathways shared between the states (Table 2). The significantly enriched cancer pathways common for all states included cell-cell communication, interactions with extracellular matrix, collagen biosynthesis, and receptor tyrosine kinase cascades (Table S7).

**Table 2.** Unique and common Reactome cancer pathways enriched in the evolutionary states.

| Unique pathways<sup>a</sup> | Common pathways<sup>b</sup> |
| --- | --- |
| Evolving:<br>• MAPK family signaling cascades<br>• Signaling by NOTCH1 | **Cell-Cell communication**<br>• Cell-cell junction organization |
| Maintaining:<br>• PI3K/AKT Signaling in Cancer<br>• VEGFA-VEGFR2 Pathway | **Adherens junctions' interactions**<br>• Extracellular matrix organization<br>• Non-integrin membrane-ECM interactions |
| Adaptive<br>• MET activates PTK2 signaling<br>• Signaling by ERBB4 | **Collagen biosynthesis and modifying enzymes**<br>• Collagen formation & chain trimerization<br>**Signaling by Receptor Tyrosine Kinases**<br>• DAG and IP3 signaling<br>• Rho GTPase cycle<br>• Signaling by VEGF |

ECM, extracellular matrix.
a Listed are pathways that were statistically significantly enriched in the state at the threshold of FDR-adjusted $p$ value $<0.05$ .
b Pathways indicated in bold were common for three states, whereas the normal font indicates daughter pathways shared by two states.

The significantly enriched cancer pathways unique to each state comprised MAPK and NOTCH1 (evolving state), PI3K/AKT and VEGFA (maintaining state), and ERBB4 and MET (adaptive state) signaling cascades (FDR-corrected *p* < 0.05). At the patient level, 49%, 28% and 23% of the patients had enriched PI3K/AKT, ERBB4, and MAPK & NOTCH pathways, respectively (Methods; Table S8).

To examine how genomic alterations affect the activity of the evolutionary state specific six signaling cascades, we obtained pathway enrichment scores per patient (*n* = 52) from 167 RNA-seq diagnostic samples matching the whole-genome sequenced samples used in the evolution analysis (Methods; Table S8). Consistent with the genomics findings, both MAPK and NOTCH1 pathways exhibited significant downregulation in tumors occupying the evolving state, VEGFA and PI3K demonstrated upregulation predominantly in tumors corresponding to the maintaining state, and ERBB4 and MET showed upregulation in tumors from mostly, but inconsistently for the MET cascade, of the adaptive state (Figure 4).

**Figure 4.**
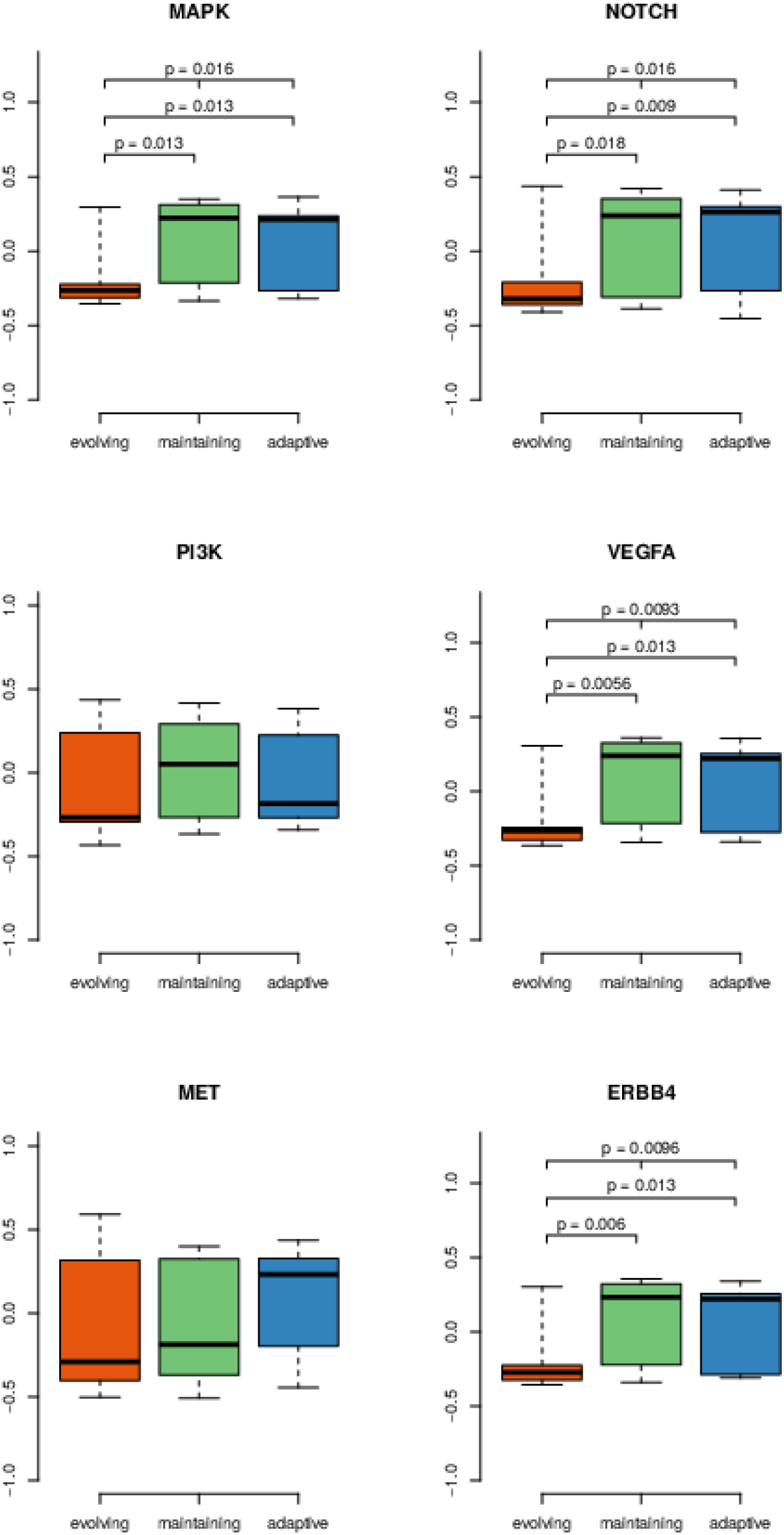
Pathway activity scores for unique signaling cascades estimated from the RNA-Seq data. Box-whisker plots of the pathway enrichment scores in RNA expression data across the evolutionary states (ANOVA, shown are *p* values < 0.05).

We next hypothesized that the differences in transcriptional profiles between the states are reflected in the cell composition. To investigate cancer, immune, and stromal components, we performed RNA decomposition (Häkkinen et al., 2021) and obtained cell compositions (Methods; Table S9). The tumors diagnosed at the adaptive state were characterized by the highest number of cancer cells and the smallest number of stromal cells, whereas immune component exhibited equally low numbers in all three states (Figure 5A).

**Figure 5.**
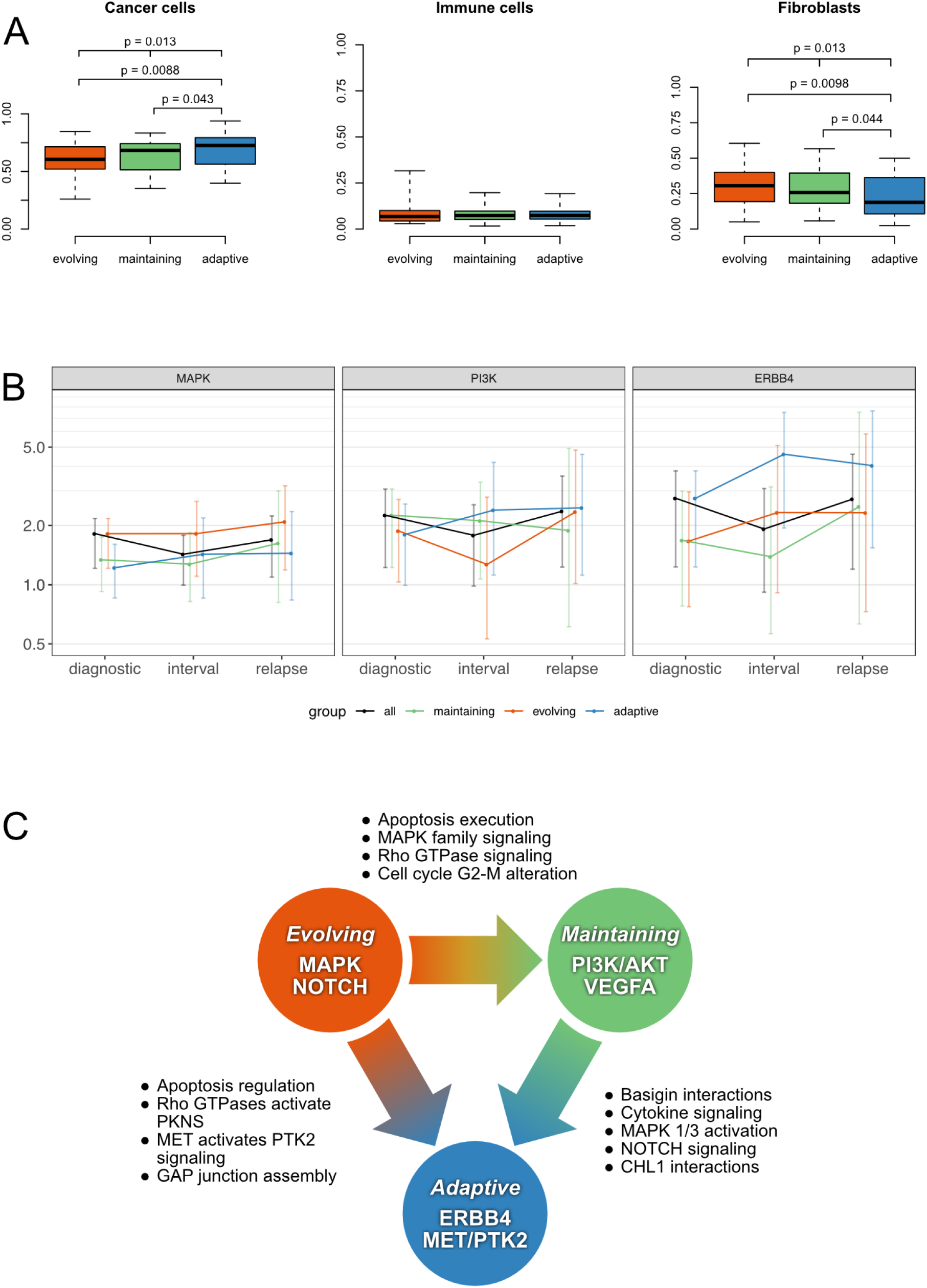
Evolutionary trajectories and stability of unique signaling cascades across the treatment. **A**. Box-whisker plots of the cell type abundances derived from the bulk RNA decomposition across the evolutionary states (ANOVA, adjusted for the tumor sites, shown are *p* values < 0.05). **B**. Enrichment of MAPK, PI3K, and ERBB4 signaling cascades at diagnosis, interval, and relapse treatment phases. The Y-axis denotes odd ratios obtained from the pathway analysis and the whiskers denote range of non-significance at *p* = 0.05. **C.** Evolutionary trajectory for three states characterized by unique signaling cascades (in nodes) and transition pathways identified by the nested pathway analysis (in clear boxes).

### MAPK, PI3K/AKT, and ERBB4 pathways are not affected by chemotherapy and remain enriched at relapse

We next assessed the effect of NACT on the evolutionary states and pathways using genomics data from 41 samples from 19 patients treated with NACT. This analysis indicated that the evolutionary state remained the same after the chemotherapy in 58% of the patients (Table S6). While clonal composition in general was stable during the treatment, the stability of the evolutionary states differed dramatically. Evolving state showed the highest instability with all tumors changing to the adaptive state after NACT. The patients from the adaptive state showed changes both to evolving and maintaining states. The maintaining state was the least prone to change, with a statistically significant lack of association with the other states (Table S6, *p* = 0.01).

Stable clonal composition of the maintaining state samples during therapy suggests that the treatment-naïve tumors contain subclones with aberrations in the PI3K/AKT signaling pathway, a known driver of tumor progression and resistance to chemotherapy (Liu et al., 2020; Masoodi et al., 2020). Conversely, tumors occupying evolving state increased both in complexity and divergence during therapy resulting in the emergence of new subclones. Among the adaptive tumors, only a fraction underwent selection resulting in the state with a lower clonal divergence after chemotherapy, such as patient EOC26 (Supplementary Jellyfish Plots).

We analyzed the stability of the six signaling cascades from 22 relapse samples from 14 patients. In the tumors in the evolving state, MAPK pathway, which was significantly enriched at treatment-naïve samples, was also significantly enriched at NACT and relapse samples. In the maintaining state, PI3K/AKT pathway remained active after NACT and was significantly enriched at relapse. The adaptive state was distinguished by significant enrichment of ERBB4 pathway at all time points (Figure 5B; Table S10; Figure S11).

### Nested pathway analysis uncovers two main evolutionary trajectories

We next hypothesized that the identified states do not exist independently but form an evolutionary path flowing from evolving state towards maintaining and adaptive states. To investigate the relatedness and transitions occurring from one state to another, we developed a nested pathway analysis approach, which maps the phylogenetic relationships between the evolutionary states (Methods). Based on the nested pathway analysis, we identified two distinct trajectories characterized by specific pathways (Figure 5C; Figure S12).

In the first trajectory, tumors from patients of evolving state progress to the maintaining state, driven by MAPK and Rho GTPases pathways, and the altered G2-M transition in the cell cycle. The second step in this trajectory is progression to adaptive state, and this transition is marked by the activation of cytokine signaling, mostly via interferons and various interleukin families, as well as by CHL1 interactions and NOTCH signaling (Table S13).

The phylogenetic relationships between the evolutionary states also suggested an alternative trajectory, where tumors occupying evolving state would directly evolve to the adaptive state, bypassing the maintaining state. This trajectory is characterized by the activation of PTK2 signaling via MET, changes in apoptosis regulation and cell communication.

## DISCUSSION

We integrated clonal composition and topology of HGSC tumors and discovered three evolutionary states organized in the order of their development. Importantly, our stratification of patients into three states prior to treatment enabled us to indicate a group of patients with tumors in the maintaining state showing fastest times to relapse. Furthermore, patients with tumors in the evolving state do not exhibit fast relapses and demonstrate most favorable survival odds. As for the patients with tumors occupying the adaptive state, they show moderate times to progression, but once relapsed, these patients have poor survival prognosis. The novel nested pathway analysis suggests two main evolutionary trajectories that describe transitions between the states. Patients whose tumors evolve directly to the adaptive state from the evolving state have better survival then patients whose tumors evolve to adaptive state via the maintaining state, the latter characterized by the PI3K/AKT pathway alterations.

The DECIDER cohort contains samples from several anatomical sites of a patient, which allowed comprehensively quantify heterogeneity in HGSC. Strikingly, tubo-ovarian tumors harbored 70% more unique clones than intra-abdominal tumors or ascites, which is in line with the observation that tumor diversity is acquired early in the primary ovarian tumors (Masoodi et al., 2020). Tubo-ovarian samples represent the site-of-origin and early local spreading (Labidi-Galy et al., 2017) and have been used extensively in HGSC research, such as in The Cancer Genome Atlas cohort (The Cancer Genome Atlas Research Network, 2011). Given the high number of unique clones in the tubo-ovarian samples, it is likely that the results based on exclusively tubo-ovarian samples are not reproducible in cohorts with samples from other anatomical sites. Another important observation from our heterogeneity analysis is that ascites samples have in general 14% less unique clones than the samples from other anatomical sites. Thus, cohorts with predominantly ascites samples may present an increased false negative rate.

Our results show the importance of longitudinal multiregion cohorts, such as the DECIDER cohort. To provide comprehensive view on our cohort, the genomics data are available via interactive visualization platform GenomeSpy (https://csbi.ltdk.helsinki.fi/p/lahtinen_et_al_2022/). GenomeSpy allows easy and fast exploration of mutations or copy-number aberrations for any gene or genome segment in the human genome in our cohort. All raw data are available in the European Genome-phenome Archive (EGA). In addition to interactive visualization of the genomics data, we developed Jellyfish plots that enable visualization of the tumor evolution progress in longitudinal, multi-sample cohorts.

The pathway analysis results suggest that MAPK and NOTCH1 are involved in cancer progression at evolving state, followed by enriched PI3K/AKT, ERBB4 and MET pathways at maintaining and adaptive states. This implies that patients stratified by the evolutionary state at the time of diagnosis could benefit from the drugs targeting these pathways, many of which were already approved or being investigated in clinical trials (Bonello et al., 2018; Diaz-Padilla et al., 2015; El-Gamal et al., 2021; Konstantinopoulos et al., 2021; Vergote et al., 2020). Importantly, MAPK, PI3K/AKT and ERBB4 pathways remain unaffected by platinum-taxane chemotherapy and are active also in relapses. This shows that the standard-of-care in HGSC is not able to eliminate the influence of these well-established cancer growth and resistance driving pathways, and combinatorial interventions are required. For example, in line with earlier studies (Dahl Steffensen et al., 2008; Davies et al., 2014; Saglam et al., 2017), activated ERBB4 pathway is enriched in one-third of patients in the DECIDER cohort and manifests a genomic landscape unique to the adaptive state. This, combined with the encouraging preclinical results for HER4-targeted inhibitors (Canfield et al., 2015; Engelman et al., 2007; Hickinson et al., 2010; H. Lee et al., 2020), supports further trials with HGSC patients. Similarly, half of the patients in the DECIDER cohort have enriched PI3K/AKT pathway. Given that PI3K/AKT was found to be active in the maintaining state tumors and in relapse samples, our results suggest combining PI3K/AKT inhibitors with drugs targeting the VEGFA-VEGFR2 pathway, which is also enriched in the maintaining state tumors, in the platinum resistant HGSC patients with maintaining state cancer and therefore the shortest survival expectation.

We assessed the effect of neo-adjuvant chemotherapy and further lines of treatment on the stability of the states and underlying unique signaling cascades. The evolutionary states remained stable in most patients across the NACT treatment. Given that ~80% of the HGSC patients respond well to the primary treatment but suffer from difficult-to-treat relapses, our approach allows identifying pathways that contribute HGSC evolution and promote platinum resistance leading to a poor survival. As resistance to platinum-based chemotherapy is a major clinical problem in the treatment of HGSC, our study expands understanding of evolutionary processes rooting chemoresistance before diagnosis.

The main limitation of this study is that the average whole-genome sequencing coverage for HGSC samples was 36x, which limits resolution to detect rare/small subclones in a sample. As our cohort contains multiple samples from the same patients, we are able to detect major clones robustly as evident from agreement of our genomic landscape results with the other HGSC cohorts with a deeper sequencing coverage. The number of samples per patient varied in our cohort, which affects the detection of clusters by PyClone VI. We tested statistically that the varying number of samples per patient does not bias our results. While we focused on tumor evolution based on data from cancer cells without explicitly integrating non-genomic information, which is suggested to affect tumor evolution (Pich et al., 2022; Zhang et al., 2018), our results from the decomposed transcriptomics data support the contributing role of stromal cells to tumor evolution, warranting future studies with histopathological and single-cell data.

In summary, our prospective, longitudinal, multiregion study allowed, for the first time, quantification of heterogeneity and clonal information from >50 HGSC patients. We identified three evolutionary states on a landscape of clonal complexity and divergence, that are characterized by distinct pathways, which provided rational targets for intervention at relapse. We showed that genomic heterogeneity in HGSC tumors varies between primary and metastatic tumor sites but, more importantly, was not significantly affected by the current chemotherapy treatment. This evidences that the current interventions curb cancer growth and tumor load rather than eradicate the cancer growth root causes leading to chemotherapy resistant disease. Our results pinpoint targets for interventions that have a higher likelihood of being effective in the treatment of relapsed HGSC patients.

## ACKNOWLEDGMENTS

This project received funding from the European Union’s Horizon 2020 Research and Innovation Programme under Grant agreement no. 965193 (DECIDER) and no. 667403 (HERCULES), the Academy of Finland (projects nos. 325956 and 322927), the Sigrid Jusélius Foundation and the Cancer Foundation Finland. We are grateful to Dr. Ann-Christin Ostwaldt for proof-reading the manuscript and Dr. Anna Vähärautio for critical comments of the manuscript. The authors wish to acknowledge the CSC-IT Center for Science, Finland, for computational resources.

## AUTHOR CONTRIBUTIONS

Conceptualization: A.L., K.L., A.V., A.H., J.O., and S.Ha. Methodology and Software: K.L., Y.L., and A.H. Formal Analysis: A.L., K.L., Y.L., S.J., A.H., and J.O. Investigation: A.L., K.L., Y.L., K.H., A.H., J.O., and S.Ha. Resources: K.L, Y.L., S.J., K.H., O.C., S.H., A.H., J.H., and S.Ha. Data curation: Y.L., A.V., J.H., J.O., and S.Ha. Visualization: K.L., A.H., and J.O. Supervison: A.H., J.O., and S.Ha. Project Administration: J.O. and S.Ha. Funding Acquisition: O.C., A.H., J.H., and S.Ha. Writing - Original Draft: A.L., J.O., and S.Ha. Writing - Review & Editing: A.L., K.L., Y.L., A.V., S.J., K.H., O.C., S.H., A.H., J.H., J.O., and S.Ha.

## METHODS

### Resource availability

#### Lead contact

Further information and requests for resources and reagents should be directed to and will be fulfilled by the lead contact, Jaana Oikkonen and Sampsa Hautaniemi.

#### Materials availability

This study did not generate new unique reagents.

#### Data and code availability

All raw DNA sequencing data will be submitted to the European Genome-phenome Archive (EGA) and will be publicly available by the time of publishing (under data access committee EGAC00001001760). Raw bulk RNA sequencing data are deposited in the EGA and are publicly available (EGAS00001004714). The patient clinical annotations are found in Table S2.

Code will be made publicly available from Github by the time of publishing.

Any additional information required to reanalyze the data reported in this paper is available from the lead contact upon request.

## Experimental model and subject details

### HGSC patients and samples

#### Cohort description

HGSC cohort comprised 55 female patients selected from the DECIDER cohort (NCT04846933), and treated at Turku University Hospital, Finland, from March 2014 to May 2019. The treatment was either primary debulking surgery (PDS), followed by a median of six cycles of platinum-taxane chemotherapy, or neoadjuvant chemotherapy (NACT), where primary laparoscopic operation with diagnostic tumor sampling was followed by a median of three cycles of NACT including carboplatin and paclitaxel. In NACT-treated patients, interval debulking surgery aiming at complete cytoreduction was performed after NACT. All patients participating in the study gave their informed consent, and the study was approved by the Ethics Committee of the Hospital District of Southwest Finland.

#### Sample selection

For this study we used 207 samples from fresh-frozen solid tumor tissues and ascites. Tumor tissues included tubo-ovarian (ovaries and fallopian tubes), intra-abdominal (omentum, peritoneum, bowel mesenterium), other sites (lymph nodes, liver), and ascites. The treatment phases represented diagnostic samples collected prior to treatment, interval samples obtained after NACT, and relapse samples collected at progression. All patients had at least two samples with estimated purity > 8% from either different treatment phases or different tissues sites. As some downstream analyses expect each patient to be represented by exactly one sample, we chose one fresh-frozen sample for each patient using the following prioritization: 1) Treatment-naïve metastasis, 2) Treated metastasis, 3) Treatment-naïve tubo-ovarian, 4) Treated tubo-ovarian, 5) Ascites. If multiple choices were available, e.g., a patient had two treatment-naïve metastasis samples, the one with the highest purity was chosen. The representative samples are indicated by a metadata attribute in the GenomeSpy visualization.

#### Patient-derived cell lines

Primary patient-derived cell cultures (n=7) were established in spheroidal DMEM-F12 medium (Thermo Fisher Scientific, USA), as described in (Kaipio et al., 2020), and were sampled for sequencing. The cultured cells were characterized by next-generation sequencing and verified to be tumorous based on a known *TP53*-mutation.

### Method details

#### Sample preparation

Tumor tissue samples were obtained from laparoscopy and debulking surgery during normal clinical treatment course and were histologically examined by a pathologist. Samples from ovaries and omentum were preferred for diagnostic and interval phases. We used ascites samples when available, especially for the relapses. For germline variation detection we collected blood samples at the beginning of treatment, and DNA was extracted in Auria biobank using Chemagic DNA Blood Kit Special (PerkinElmer Inc., USA) and Chemagic 360 instrument (PerkinElmer Inc., USA). For other samples, Qiagen AllPrep kit was used to extract DNA and RNA simultaneously.

#### Sequencing

Tissue samples with sufficient DNA/RNA content and quality were sent to library preparation and sequencing to BGI (BGI Europe A/S, Denmark), where whole-genome sequencing (WGS) was performed with either DNBSEQ (BGISEQ-500 or MGISEQ-2000, MGI Tech Co., Ltd., China) or HiSeq X Ten (Illumina, USA) as 150bp paired end sequencing. Median coverage was ~36x (range 24-96x). RNA sequencing was performed with DNBSEQ, HiSeq X Ten or Illumina Hiseq 4000 (Illumina, USA) as 100bp or 150bp paired end sequencing.

### WGS data preprocessing

We performed WGS analyses using Anduril, a bioinformatics workflow platform developed for large data sets (Cervera et al., 2019). We trimmed read data using using Trimmomatic 0.32 (Bolger et al., 2014) and assessed quality with FastQC v0.11.4 (Andrews, 2015) via Anduril QCFasta v5.0 component. The trimmed reads we aligned using BWA-MEM 0.7.12-41039 (Li, 2013) with option -M against the human reference genome GRCh38.d1.vd1, which was followed by read deduplication using Picard 2.6 (https://github.com/broadinstitute/picard) *DuplicateMarker* and base quality score recalibration using Genome Analysis Toolkit (GATK) (McKenna et al., 2010) version 3.7 *BaseRecalibrator* (DePristo et al., 2011).

In one batch we filtered vector contamination by aligning first the trimmed reads against the vector sequence, keeping only unmapped reads which were converted back to FASTQ using BEDtools 2.28.0 (Quinlan, 2014; Quinlan & Hall, 2010) *bamtofastq* before proceeding with the alignment as above. We used the following vector sequence for read filtering: AACAATTTCACACAGGAAACAGCTATGACCATGATTACGCCAAGCTCGAAATTAACCCTCACTAAAGGGAACAAAAGCTGGAGCTCCACCGCGGTGGCGGCCGCTCTAGAACTAGTGGATCCCCCGGGCTGCAGGAATTCGAT.

Finally, we computed alignment stats using Anduril BamStats component which includes coverage estimation via BEDtools *genomecov*. We also estimated cross-sample contamination using GATK 4.1.4.1 according to the best practices (van der Auwera & O’Connor, 2020). The primary normal samples of each patient used in variant calling served as the matched normals in the contamination estimation. We filtered samples with contamination greater than 10%.

### Mutation calling in WGS data

#### Panel of normals

We used a somatic panel of normals (PoN) generated from our blood derived normal samples in somatic variant calling according to GATK best practices. More specifically, we used primary normal samples of each patient with contamination less than 5%, totaling 80 DECIDER normals. Furthermore, as germline resource we used Finnish allele fractions from gnomAD v3.0 (Karczewski et al., 2020), including passing variants as well as those only failing the filter ‘InbreedingCoeff.

#### Somatic variants

We called somatic short variants by jointly calling multiple tumor samples against a single matched normal for each patient using GATK *Mutect2* based on best practices. As in PoN, we utilized Finnish gnomAD allele frequencies as germline resource. Variant calling was restricted to primary chromosome assemblies excluding Y chromosome. We filtered the variants using GATK *FilterMutectCalls*, keeping only variants passing all filters. Since somatic calling preceded purity estimation, our per-patient joint calling sample set contained low purity samples excluded from this study. We kept variants that had ALT reads in the retained samples. In variant annotation we used GATK *VariantAnnotator* to add dbSNP 153 (Kitts et al., 2013; Sherry et al., 2001) IDs and utilized the offline version of Combined Annotation Dependent Depletion (CADD) v1.6 (Rentzsch et al., 2021) with an in-house solution to write into variant call format files. We computed variant allele frequencies (VAF) from reference and alternate (ALT) allele read depth (AD) field. Finally, we used ANNOVAR 20191024 (K. Wang et al., 2010) to include the following annotations: refGene (Pruitt et al., 2014) dated 2020-03-01, downloaded from the University of California, Santa Cruz (UCSC) genome browser database (Navarro Gonzalez et al., 2021) and available online at https://hgdownload.soe.ucsc.edu/goldenPath/hg38/database/, as well as other annotations converted for ANNOVAR including ClinVar 20200506 (Landrum et al., 2018), customized annotations for COSMIC v91 (Tate et al., 2019), and gnomAD v3.0 genomes.

#### Germline variants

We called germline short variants using GATK joint genotyping approach following best practices. More specifically, we used GATK 4.1.4.1 *HaplotypeCaller* to call in genomic variant call format (GVCF) on individual normal samples from a set of 80 DECIDER normal samples with allele-specific annotations before merging and genotyping with *GenotypeGVCFs*. We restricted variant calling to primary nuclear chromosome assemblies excluding Y chromosome. In variant allele filtering, we used allele-specific variant quality score recalibration, dropping all filtered ALT alleles from subsequent analyses.

We further refined the genotype calls using GATK *CalculateGenotypePosteriors* and filtered genotypes with genotype quality (GQ) less than 20 and read depth (DP) less than 10. We then filtered dubious genotype calls by requiring the sum of depths for genotyped allele reads be a minimum of 5 and VAF of genotyped alleles be at least 20% (AlleleBalance filter), both computed from the AD field.

### CNV calling from WGS data

We used the GATK (McKenna et al., 2010) version 4.1.4.1 to perform the copy-number segmentation. To collect the minor allele counts (BAF), we used all filtered biallelic germline SNPs with heterozygous calls (VAF between 40% and 60%) from each patient. Read-count collection used one kilobase intervals. Both read and allelic count collection excluded regions listed in the ENCODE blacklist (Amemiya et al., 2019a). To denoise the read counts, we used platform specific (HiSeq, DNBSEQ) panels of normals built from the normal samples. After the segmentation, we used ASCAT v2.5.2 (van Loo et al., 2010) to estimate purity, ploidy, and allele-specific copy numbers. Since ASCAT does not readily accept pre-segmented data, we converted the segmented data into “faked” SNPs readable by ASCAT. We also used VAFs of the *TP53* mutations as additional evidence for the optimal ploidy/purity selection. Since all the patients had homozygous *TP53* mutations in their cancer cells, we could use the VAF and total copy number (CN) to calculate the purity:

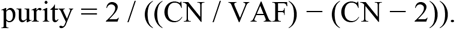

When ASCAT and *TP53*-based purities disagreed, we chose the higher one. Samples with no abnormal CNV model found were excluded from further analyses.

As the contribution of non-aberrant cells on the logR and BAF values encumber further analyses, we calculated “purified” values, i.e., removed the non-aberrant component, using the following formulas.

Purified R, based on discussion in https://github.com/lima1/PureCN/issues/40:

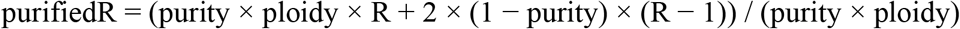

Purified BAF, derived from S2, S7, and S8 of (van Loo et al., 2010):

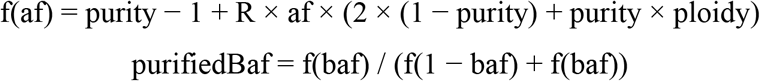

### GISTIC analysis

We ran GISTIC 2.0.23 (Mermel et al., 2011) with the default parameters for all 55 patients using the representative samples and purified logR values. To exclude poorly aligned regions and population-level copy-number variance from the analysis, we applied our internal DECIDER blacklist, which includes regions that have abs(logR) > 0.2 in at least three of the 114 available normal samples. The blacklist is available in the GenomeSpy visualization as an annotation track.

### GenomeSpy visualization

All genomics and key clinical data accompanied by bookmarks are available via an interactive visualization tool GenomeSpy (https://genomespy.app/) at https://csbi.ltdk.helsinki.fi/p/lahtinen_et_al_2022/. In brief, we specified an interactive visualization that comprises copy-number variation, loss of heterozygosity, somatic short variants (SSVs), essential metadata such as clinical variables, and GISTIC results. Only those SSVs were included that had a CADD score greater or equal to 10.0 or were pathogenic according to ClinVar. The visualization contains additional tracks that provide context for the genomic data: cytobands, ENCODE Blacklist (Amemiya et al., 2019), RefSeq Gene annotations (O’Leary et al., 2016), and COSMIC Cancer Gene Census (Sondka et al., 2018).

### Subclonal mutations

For subclonal clustering, we included only mutations that had 1) CNV segment information in all samples, 2) at least two ALT reads in a tumor sample, 3) no ALT reads in matched normal sample, 4) median coverage at least 20, and 5) minimum 15 reads in every sample of that patient. Mutations were divided into cancer cell fraction (CCF) clusters using PyClone-VI (Gillis & Roth, 2020). It corrects for copy-number status of the mutations and estimates CCF status across samples. We estimated the model for a maximum of 30 clusters and used 100 restarts. The primary run was performed using only single base substitutions (SBS). Indels (short insertions and deletions) were fitted to the clusters via a secondary run with reduced weight for the indels: indel read counts were divided by 2. Indels from secondary clusters were merged into original clusters when the clusters had at least 70% overlap in SBS. Original, SBS based cellular frequencies were used in downstream analyses. Indels have less reliable VAF values, and this protocol was used to build the most reliable clusters.

### Subclonal cluster filtering

We kept clusters with 5% minimum CCF in at least one sample. The founding cluster was estimated from the cellular frequencies as the most frequent one across samples. Additional clusters with higher CCF than that of the founding cluster (allowing for 5 percentage point error) were excluded. Mutations with cluster assignment probability less than 0.6 and the median probability were excluded.

For further filtering of clusters with an excess of likely artefact mutations, we employed mutational signature analysis combined with additional variant filtering. For mutational signature analysis, we generated cluster mutational spectra from their non-mitochondrial single nucleotide variants (SNVs) as well as patient spectra from their sums. We fitted signatures using the function pnnls from R package lsei (Y. Wang et al., 2020) against COSMIC v3.1 signatures corrected for GRCh38 trinucleotide frequency excluding chromosomes Y and M. Signature selection was based on SigProfilerAttribution (Alexandrov et al., 2020) with backward and forward selection evaluating cosine similarity with thresholds 0.01 and 0.05 respectively as well as connected signature and forced signatures (SBS1 and SBS5) rules. Briefly, we started with common ovarian cancer signatures SBS1, SBS2, SBS3, SBS5, SBS12, and SBS40 and fit the patient level spectra. The patient signatures together with common signatures were initial signatures for preliminary cluster signature selection, the result of which were used as inputs for the final patient level signature selection but with a less stringent backward selection threshold of 0.001. The resultant patient signatures were used as inputs for the final individual cluster signature selection of the respective patient.

As additional filters, we removed variants with NALOD less than 1.0 or CONTQ less than 10 as well as those present in an extended PoN. The extended PoN was generated from 105 DECIDER and 99 TCGA normals using GATK 4.1.9.0 and global allele frequencies from gnomAD v3.0 with maximum germline probability set to 1.0. We chose the TCGA samples from the NCI Genomic Data Commons (GDC), including the legacy repository, by selecting “TCGA” program name, “blood derived normal” sample type, and “WGS” experimental summary as common criteria. In GDC repository we further specified “illumina” platform, while in the legacy GDC, we selected those with “HiSeq X Ten” platform, “BAM” data format, and “unaligned” or “aligned” data type. We chose the samples on December 22, 2020, and in total obtained 99 normal samples from TCGA sub-studies: four SARC (sarcoma), 13 ESCA (esophageal carcinoma), and 82 LUAD (lung adenocarcinoma). The data were harmonized to our preprocessing. However, base quality recalibration was not repeated for LUAD data.

To filter clusters, we utilized the additional filters with signature fitting in three analysis combinations: runs with 1) extra forced signatures with extended filtering, 2) extra forced signatures, and 3) neither. The extra forced signatures were SBS34, SBS41, SBS47, SBS57, SBS58, and SBS90, which were common among spurious clusters. We computed the proportion of artefact mutations in clusters as the total fractional contribution of signatures SBS34, SBS41, SBS47, SBS57, SBS58, SBS90, and SBS7c. We then computed the cluster artefact score (a) as a weighted average, using cosine similarity of fit, of artefact proportion of runs 2 and 3. We also computed the proportion of likely artefacts (b) by summing the fraction of variants excluded by the additional filters and the proportional contribution of artefact signatures of run 1 scaled by the fraction of passing variants. Clusters with an artefact score (a) greater than or equal to 0.25 and proportion of likely artefacts (b) greater than or equal to 0.50 were filtered, as were clusters with fewer than 30 SNVs after variant filters of run 1.

After all filters, 427 clusters remained (77.5% of all clusters).

### Phylogenetic trees

Mutational trees for all patients were estimated from clustered mutations with ClonEvol (Dang et al., 2017). Frequency threshold of 1% was used as a positive threshold for a cluster to exist in a sample. A solution that included the maximum number of clusters was selected. When multiple options with the maximum number of clusters were available, solutions excluding smallest (least mutations) or truncal-like clusters, were favored. The truncal-like clusters were assumed to be possible errors in clustering, and they were identified by having in all samples CCF of more than half of that of the founding cluster. When multiple trees were available for same clusters, the best tree was selected using pigeon-hole principle (Gundem et al., 2015). 76% of the 427 clusters were fitted to the trees.

### Mouse contamination filter

We observed mouse contamination in three tumor samples (EOC423_pOme2_DNA1, EOC1120_pPer1_DNA1, and EOC1129_pPer2_DNA1) affecting somatic calling results, which resulted in two of these samples having a spurious cluster each. The two clusters were removed as a result. To minimize the effect of contamination in other downstream analyses, we realigned the corresponding BAMs against a concatenated reference genome of GRCh38.d1.vd1 and mouse genome build GRCm39 (toplevel) where the mouse contigs served as decoys. We also reran duplication marking. With the realigned BAMs, we force-called mutations private to the contaminated samples using *Mutect2* in the affected patients, otherwise with the same parameters as in the original somatic variant calling. We tagged variants that no longer had ALT reads in the realigned samples and removed them in downstream analyses.

### Jellyfish plots

Phylogeny and clonal compositions estimated by ClonEvol served as input for the patients’ jellyfishes. The visualizations are largely hand-drawn using Affinity Designer (RRID:SCR_016952). However, we extracted the clonal compositions from the ClonEvol data structures and used depth-first search on the phylogenetic tree to find a consistent order for the subclones. Subsequently, the clonal compositions were plotted as stacked bar charts using R package ggplot2 and exported as Scalable Vector Graphics format for manual editing and annotation.

### Frequently mutated genes

We analyzed frequently somatically mutated genes using dNdScv (Martincorena et al., 2017) to correct the background mutation rate of each gene. Since the tool requires mutations to be independent, we used patient representative samples (N=55). Furthermore, phased pairs of variants from *Mutect2* with less than 10 bases between the last affected base of the first variant and the position of the next variant were merged as complex substitutions. We ran dNdScv with Y chromosome genes excluded and using GRCh38 covariates and RefCDS inputs downloaded from https://github.com/im3sanger/dndscv_data/tree/master/data on January 27, 2022. Following dNdScv’s results, we report patients with missense, nonsense, splice site or complex mutations (incl. indels and multiple nucleotide variants), omitting stop loss SNVs.

### Subclonal heterogeneity

We used clonal information obtained via mutation trees of 207 tumor tissue samples from all patients to estimate clonal complexity and divergence.

#### Clonal complexity

To measure clonal complexity of each individual sample, we quantified the perplexity of the subclonal cancer cell frequency distribution of the corresponding mutation tree. The perplexity is an estimate for the effective number of clones, by taking their cellular frequencies into account and is immune to the number of called clones (e.g., *C* = *k*’ for *k*’ equifrequent clones regardless of *k*). The clonal complexity of the *j*^th^ sample is computed as follows:

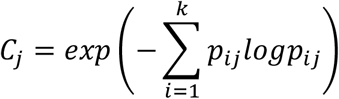

where *p_ij_* is the normalized frequency of the *i*^th^ clone over the *k* clones of the *j*^th^ sample, and by definition 0 · *log*(0) = 0. The clonal complexities were averaged in entropic (logarithmic) space to capture patient, tissue, and phase (diagnostic-interval) specific clonal complexity. We also evaluated patient-tissue specificity within patients and tissues using the logarithmic complexity values. This analysis reveals which tissue sites exhibit heterogeneity within the sample, not across the patient disease.

#### Clonal divergence

We also measured whether each sample’s clonal frequency distribution differs from the others. This was quantified using the Kullback-Leibler divergence of the sample’s clonal distribution from the average distribution over all samples of each individual patient. The corresponding perplexity represents the effective number of unique clone changes to a particular sample compared with an average sample of the patient. The clonal divergence of the individual *j*^th^ sample is computed as follows:

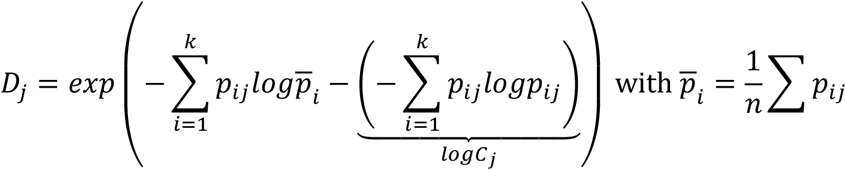

where *p_ij_* is the normalized frequency of the *i*^th^ clone over the *k* clones of the *j*^th^ sample, 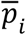 is the average frequency of *i*^th^ clone in the reference (the *n* samples of the patient), and 0 · *log*(0) = 0. The clonal divergencies were averaged in entropic (logarithmic) space to determine the global average effect for different tissues sites and their significance.

#### Statistical analysis

We applied variance analysis (ANOVA) to assess if the patients, tissues, and phase components feature distinct, specific degree of heterogeneity in their tumors for both the clonal complexity and divergence. First, we assessed if the patient specific complexity is significant versus the unexplained complexity (the replicate variation excluding patient, tissue, and phase specific variation in these data). We performed a similar test for tissue specific and treatment phase specific complexity versus unexplained complexity. We also tested whether the tissue sites exhibit heterogeneity within the patient specific complexity and vice versa, quantifying changes in specific but distinct tissue sites or patient groups. Similarly, we performed ANOVA of the cross-entropy heterogeneity to determine which factors affect the corresponding clonal divergence. The ANOVA coefficients correspond to logarithmic effective clonal number ratios, i.e., a negative coefficient indicates a decrease and positive increase in the effective number of intratumor or tumor-unique clones.

### Clustering patients by heterogeneity measures

A patient stratification was obtained by clustering the patients in the clonal complexity versus clonal divergence space. The clonal complexity 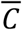 and divergence 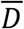 of a patient is the geometric mean of the sample complexities *C_j_* and divergencies *D_j_* of the patient (arithmetic entropic mean), respectively. In particular, when averaged over all the samples, the clonal divergence 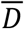 represents the effective number of unique clone sets within the set of samples of a patient.

Since the clonal divergence 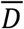 can only be unity when a single sample is available from the patient, a weighted K-means clustering approach with object-by-feature specific weights was used (Cordeiro de Amorim & Mirkin, 2012). Specifically, the following distance metric *d* is implied:

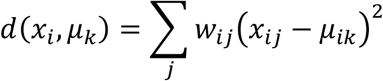

giving optimal cluster centroids of:

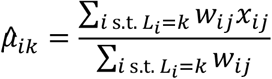

of weighted averages of the data, where *x_ij_* is the *j*:th component of the *i*:th sample, *w_ij_* is the corresponding weight, *μ_k_* is the centroid of the *k*:th cluster, and *L_i_* is the label of the *i*:th sample. The weights are set to zero for missing data and are unity otherwise, implying the missing data have no influence on the clustering along the missing dimension, but can affect the clustering along the others. For visualization purposes, the missing values were imputed with the most likely values (i.e., minimal distance), that is, the corresponding cluster centroid. Note that the imputation occurs post-hoc and does not affect the clustering.

The clonal complexity and divergence were computed for the diagnostic, interval, and relapse samples separately, and the data from the diagnostic samples was used for patient stratification. For the clustering, 1,000 iterations and 1,000 restarts were used. Among the 2D discriminators, *K* = 3 clusters was favored by a series of likelihood ratio tests, which was also the preferred model according to the Akaike Information Criterion (Yang, 2005).

### Estimation of clonal pseudo-age

To quantify the clonal topology, we computed the branch and mutation depth and diameter. The branch depth and diameter are computed from the graph induced by the clonal subdivision tree with unit edge lengths, while for the mutational depth edge lengths with a difference in mutations is used.

The depth is defined as the clonal frequency weighted average distance from the root to the observed clones, while the diameter is the weighted average distance within the observed clones. Specifically, given *D_ij_* is the shortest undirected path length from the *i*:th to the *j*:th clone and *p_i_* is the observed clonal frequency, the depth is given by:

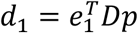

where *e_1_* = (1,0,…) is frequency distribution of the root (*i* = 1) and the diameter by:

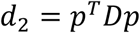

The branch depth acts as the average pseudo-age in number of clonal subdivisions, the diameter measuring its variance, while the mutational depth and diameter are the corresponding measures of a pseudo-age in the number of mutations. The branch and mutational depth and diameter agreed qualitatively, but we found that the branch depth more accurately captured the clonal tree topology between the samples.

### Pathway analyses at the subclonal level

#### Pathways enriched in the evolutionary states

To explore the biological pathways enriched at the subclonal level in the evolutionary states, we combined branch mutations for all patients included in each of the evolutionary cluster and annotated the corresponding genes (2,349, 2,363, and 2,699 genes for ‘evolving’, ‘maintaining’, or ‘adaptive’ clusters, respectively). The mutations were filtered using CADD score 10, excluding intergenic and benign mutations from ClinVar functional annotation (Landrum et al., 2018), with only branching mutations (‘non-truncal’ clusters in the downstream analysis) selected for further analyses. Pathway analysis was performed using Reactome Knowledgebase (Jassal et al., 2019) for selected gene sets and statistical significance of results was defined using a threshold of *p* value < 0.05 (after Benjamini-Hochberg adjustment). Shared pathways were defined as pathways statistically significantly enriched in any two or in all three states. Six unique signaling cascades were defined as cancer pathways a) statistically significant in one of the states and b) including known anticancer drug targets. Analyses were conducted in R software v3.6.3 (https://www.r-project.org/), using R package ReactomePA v1.38.0 (Yu & He, 2016).

#### Pathways enriched at the level of patients

To explore the biological pathways at the level of individual patient, we used branch mutations detected per patient and annotated the corresponding genes. The filtering of mutations and pathway analyses were conducted in the same way, as described for the evolutionary states above. The statistical significance of results was assessed using a threshold of unadjusted *p* value ≤ 0.1.

#### Stability of selected signaling cascades across treatment

The stability of six unique signaling cascades across the treatment was assessed in the samples collected at diagnosis, interval, and relapse phases (149, 43, and 22 samples, respectively). The filtering of mutations and pathway analyses were conducted in the same way, as described above, and statistical significance of results was defined using a threshold of *p* value < 0.05 (after Benjamini-Hochberg adjustment).

### Nested pathway analyses

A typical binary enrichment analysis, e.g., of abundance of pathway specific hits, uses the overlap count of the called genes and the genes involved in the pathway and a Fisher’s exact test to derive significance (Yu & He, 2016) for over- or underrepresentation under the assumption that the two variables are unrelated. A corresponding test for an increasing (or decreasing) enrichment substitutes the Fisher’s noncentral hypergeometric distribution for the null model (Liao & Rosen, 2001) in place of the centered hypergeometric distribution.

By assuming an evolutionary directed acyclic graph (not to be confused with the clonal tree graph) of groups that retains the degree of previous acquired pathway aberrations and introduces novel ones, a series of noncentral hypergeometric tests can be constructed to select a model among the candidate evolutionary graphs. Given the groups evolving (E), maintaining (M), adaptive (A) and their determined timing, the candidates are: E M A, E M>A, E>A M, E>A M>A, E>M A, E>M M>A, E>M E>A, E>M E>A M>A, where A>B represents an edge and A B no edge in the evolutionary graph.

Specifically, the maximum likelihood (ML) estimates of the noncentral hypergeometric models are first fit for all the partitions of the nodes of the evolutionary graph. In our setting, this involves the patient groups E, M, and A, giving the partitions EMA, E|MA, EM|A, EA|M, and E|M|A. Here, AB implies that the nodes A and B belong to the same subset of a partition and share the enrichment odds, while A|B implies two partitions where the odds can differ. Given a partition *P* with labeling *L*., and a set of odds ratios *ω_k_*, one for the *k*:th subset *P_k_* of the partition each, the likelihood is given by:

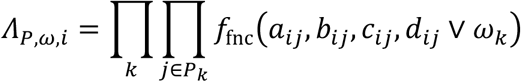

where (*a_ij_*, *b_ij_*; *c_ij_*, *d_ij_*) are the counts of the 2-by-2 crosstabulation regarding the *i*:th test (e.g. the *i*:th pathway) on the *j*:th node and *f_fnc_* is the density of the noncentral hypergeometric distribution. In practice, the maximum likelihood estimate 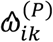 for each test and partition can be found numerically using sectioning.

The corresponding ML fits constrained by an evolutionary graph can be acquired from the above fits, namely, they occur at the ML estimates of one of the partitions whose ML estimates satisfies the evolutionary enrichment constraints as discussed above. Specifically, we write the likelihood of an evolutionary graph *G* as:

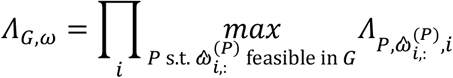

where a set of parameters *ω* are feasible if *(ω_i,L_p__* ≥ 1 ∧*ω_i,L_q__* ≥ *ω_i,L_p__*) ∨ (*ω_i,L_p__* ≤ 1 ∧ *ω_i,L_q__* ≤ *ω_i,L_p__*) for all edges from *p* to *q* in *G*, that is increased or decreased odds become no less extreme along the evolutionary graph. The appropriate evolutionary graph is selected using a series of likelihood ratio test comparing for nested evolutionary candidate graphs (cf. table at Figure S12) with degrees of freedom equal to the total number of required partitions. The disconnected graph “E M A”, where the groups are independent with no constraints, is always a putative candidate (cf. Figure S12), however, more constrained models should be favored if the evidence permits.

In a similar fashion, a likelihood ratio can be used to assess whether a specific pathway increases or decreases in enrichment along an edge from the *p*:th to *q*:th node of the selected evolutionary graph by comparing the model *Ĝ* with one with the appropriate edge deleted *Ĝ* \ (*p, q*. The pathway edge enrichment tests for the selected model E>M E>A M>A, with one of the edges E>M, E>A, or M>A deleted tabulated in Table S13.

### RNA–seq data analyses

The preprocessing and subsequent decomposition of 167 RNA-Seq diagnostic samples was performed as described in (Häkkinen et al., 2021). Pathway enrichment scores in sample and patient level for six unique signaling cascades were computed with 20,000 permutations using R package fgsea. First, we ranked each gene across all the samples, and normalized the rank values to be in [0,1]. Then, for each sample, we ordered the rank values inside out and in a decreasing order as inputs in fgsea analysis. The enrichment scores and adjusted *p* values are obtained for each individual sample. To get the patient level enrichment scores, we summed up the RNA expression data for each patient (with multiple samples), and then used that as the input for the ranking and ordering procedure. The same fgsea analysis was conducted in patient level and the corresponding enrichment scores and adjusted *p* values were obtained here as well. Cell type composition was quantified using PRISM (Häkkinen et al., 2021). The downstream statistical analyses of the enrichment scores and cell type abundances (excluding samples from ascites) were performed using ANOVA, and statistical significance of results was defined using a threshold of *p* value < 0.05.

## Quantification and statistical analysis

All statistical details of the analyses are defined in the corresponding method section.

## Additional resources

This study is a part of DECIDER trial, ClinicalTrials.gov Identifier: NCT04846933, accessed at https://clinicaltrials.gov/ct2/show/NCT04846933.

The visualization of genomic data was performed using GenomeSpy (https://genomespy.app/) and is available at https://csbi.ltdk.helsinki.fi/p/lahtinen_et_al_2022/.

